# Conserved perception of host and non-host signals via the a-pheromone receptor Ste3 in *Colletotrichum graminicola*

**DOI:** 10.1101/2024.07.27.605416

**Authors:** Anina Yasmin Rudolph, Carolin Schunke, Daniela Elisabeth Nordzieke

## Abstract

Understanding the interactions between fungal plant pathogens and host roots is crucial for developing effective disease management strategies. This study investigates the molecular mechanisms underpinning the chemotropic responses of the maize anthracnose fungus *Colletotrichum graminicola* to maize root exudates. We identify the 7-transmembrane G-protein coupled receptor (GPCR) CgSte3 as a key player in sensing both plant-derived class III peroxidases and diterpenoids. Activation of CgSte3 initiates signaling through the Cell Wall Integrity Mitogen-Activated Protein Kinase (CWI MAPK) pathway, facilitating the pathogen’s growth towards plant defense molecules. The NADPH oxidase CgNox2 is crucial for peroxidase sensing but not for diterpenoid detection. These findings reveal that CgSte3 and CWI MAPK pathways are central to *C. graminicola’s* ability to hijack plant defense signals, highlighting potential targets for controlling maize anthracnose.

## 1 Introduction

Root-infecting fungi pose a significant threat to food-producing plants, reducing both the yield and quality of harvested crops and fruits (Fisher et al., 2012). Understanding the mechanisms of fungal host root detection is crucial for developing sustainable crop protection strategies. The hemibiotrophic ascomycete *Colletotrichum graminicola* causes anthracnose in *Zea mays*, a disease characterized by brown lesions on leaves and stems (Bergstrom and Nicholson, 1999). The resulting yield loss is estimated at 10-20% worldwide annually for anthracnose stalk rot alone (Belisario et al., 2022). *C. graminicola* produces two types of asexual spores, oval- and falcate-shaped conidia, both capable of causing lesions on leaves (Nordzieke et al., 2019b). Oval conidia, asexual spores pinched off from hyphae in the vascular system and stored in adjacent parenchyma cells, are responsible for root infection (Belisario et al., 2022, Panaccione et al., 1989, Sukno et al., 2008). After rapid germination in soil, these oval-shaped spores sense and redirect their growth towards host root exudates (chemotropism), followed by invasion of the host via its roots (Rudolph et al., 2024). In contrast, falcate conidia, produced in asexual fruiting bodies (acervuli), fail to invade roots under natural infection conditions (Panaccione et al., 1989, Rudolph et al., 2024).

In recent years, class III peroxidases (Prx) have been identified as attractant molecules for root pathogenic *Fusarium* and *Verticillium* species (Turra et al., 2015, Vangalis et al., 2023). For successful perception of plant Prx, these enzymes are activated by H_2_O_2_ generated by the reduction of O_2_ by fungal NADPH oxidase (Nox) 2 complex (O_2_ → O_2_^−^) and extracellular superoxide dismutase (Sod) (O_2_^−^ → H_2_O_2_) (Nordzieke et al., 2019a). The activated Prx is then sensed via fungal pheromone receptors Ste2 and Ste3, first described for their role in sensing α- and a-pheromones in the yeast *S. cerevisiae* (Nordzieke et al., 2019a, Turra et al., 2015, Vangalis et al., 2023, Sridhar et al., 2020, Ramaswe et al., 2024, Nakayama et al., 1985).

The provision of cellular reactive oxygen species (ROS) by Nox complexes is a well-known phenomenon in fungal, mammalian, and plant cells, regulating reproduction, signaling, defense against harmful organisms, and pathogenicity (Sagi and Fluhr, 2006, Vermot et al., 2021, Nordzieke et al., 2019a, Ryder et al., 2013, Dirschnabel et al., 2014, Lambert and Brand, 2009). In fungi, Nox1/A and Nox2/B complexes contribute to proper development and interaction with host plants. Despite sharing a set of common regulatory proteins, their functions are highly specific and not interchangeable. Phenotypic characterization of deletion mutants revealed that Nox1/A is required for developmental processes such as germling fusion and sexual development, while the Nox2/B complex regulates the germination of sexual spores (Dirschnabel et al., 2014, Cano-Dominguez et al., 2008, Wang et al., 2014, Brun et al., 2009). Apart from Nox2’s role in host recognition processes, both Nox complexes are involved in the penetration of plant surfaces. In *Magnaporthe oryzae* and *Fusarium* species, Nox1/A and Nox2/B are required for proper septin ring formation, a prerequisite for penetration peg formation and elongation, respectively (Ryder et al., 2013, Wang et al., 2014, Nordzieke et al., 2019a). However, reports on *Botrytis cinerea* and *Colletotrichum* species suggest that Nox1/A, but not Nox2/B, is dispensable for the proper function of appressoria (Liu et al., 2022, Segmüller et al., 2008, Fu et al., 2022). Although there is experimental evidence that Nox-derived ROS functions via the regulation of actin (Ryder et al., 2013, Liu et al., 2022), the molecular processes resulting in Nox activation and induced downstream processing are not fully understood.

In contrast to the described findings, our lab recently identified maize-secreted diterpenoids, rather than Prx, as the attractant molecules responsible for inducing *C. graminicola* chemotropic growth towards maize roots. The a-pheromone receptor Ste3 of *C. graminicola* is responsible for the perception of these secondary metabolites (Rudolph et al., 2024). In this study, we elucidate conserved molecular processes determining chemotropic growth to plant root signals. We demonstrate that *C. graminicola* germlings can sense and redirect their growth to Prx signals in patterns similar to those observed in *Fusarium oxysporum* f. sp. *Lycopersici*. Investigation of a *Cgnox2* deletion mutant provides evidence that the Nox2 complex is essential for adequate leaf penetration via appressoria and hyphopodia and for chemotropic sensing of Prx. However, *Cgnox2* is dispensable for host plant-derived diterpenoid sensing and maize root infection by *C. graminicola*. Using genetic experiments, we show that the a-pheromone receptor CgSte3 and the Cell Wall Integrity (CWI) Mitogen-Activated Protein Kinase (MAPK) scaffold protein CgSo mediate the perception of diterpenoids and Prx. Our findings reveal that conserved molecular pathways for the perception of root-derived signals are shared among several plant pathogenic fungi, independent of their importance for recognizing the appropriate host.

## 2 Materials and Methods

### 2.1.1 Strains, growth conditions, and collection of spores

As wildtype strain, the sequenced *C. graminicola* (Ces.) G.W. Wilson (teleomorph *Glomerella graminicola* D. J. Politis) strain M2 was used (also referred to as M1.001) (Forgey et al., 1978, O’Connell et al., 2012). Oval and falcate conidia were cultivated and harvested as described previously (Rudolph et al., 2024).

### 2.1.2 Generation of plasmids and *C. graminicola* strains

A ΔCgnox2 deletion mutant and the complemented strain ΔCgnox2::Cgnox2 were generated. DNA hydrolysis and sequencing, using appropriate enzymes and primers, verified all plasmids. The oligonucleotides, strains, and plasmids used are listed in supplementary tables S1-S3.

For the generation of *Cgnox2* (GLRG_09327) deletion, the plasmid pCgnox2_KO was assembled using a split marker approach (Catlett et al., 2003) followed by subcloning in pJET1.2/Blunt (Thermo Fisher Scientific). 5’ and 3’ regions of *Cgnox2* were amplified using oligonucleotide pairs nox2_PF/ nox2_PR (1,149 bp) and nox2_TF/nox2_TR (1,160 bp), respectively. In a second step, those regions were fused to an inverted *hph* cassette (hpf-f/hph-r, 1,417 bp), mediating the resistance to hygromycine B, with the oligonucleotides nox2_PFN/nox2_TRN (3,436 bp). The plasmid pCgnox2_nat for was assembled using the NEBuilder HiFi DNA Assembly Cloning Kit (New England Biolabs) according to the instruction manual. 5’ and 3’ regions of *Cgnox2* were amplified together with the *Cgnox2* gene in a PCR using genomic DNA (gDNA) of *C. graminicola* as a template and the oligonucleotides nox2_P_comp_fw/nox2_T_comp_rv (4,093 bp). As the backbone for the assembly reaction served pJet_nat linearized with *Eco*RV (Nordzieke, 2022), mediating resistance to nourseothricin-dihydrogen sulfate in *C. graminicola* transformed with this plasmid.

Prior to transformation, the plasmids pCgnox2_KO and pCgnox2_nat were linearized using the enzymes *Hind*III and *Pvu*I, respectively. Oval conidia of CgM2 (transformation of pCgnox2_KO) or ΔCnox2 (transformation of pCgnox2_nat) served as the basis for the generation of protoplasts as described previously (Groth et al., 2021). After transformation, regenerating protoplasts were selected on a medium containing hygromycin B (500 µg/ ml, transformation of pCgnox2_KO) or nourseothricin-dihydrogen sulfate (70 µg/ ml, transformation of pCgnox2_nat). Single spore isolations were performed of antibiotic-resistant and PCR-verified primary transformants to obtain homokaryotic strains (Nordzieke, 2022). Single spore isolates of ΔCnox2 were verified by Southern Blot analyses. The hydrolysis of gDNA was performed with the enzyme *Bgl*II (Thermo Fisher Scientific). For visualization of successful deletion of the *Cgnox2* gene, the 3’ region of *Cgnox2* was amplified in a PCR reaction (Nox2_probe_F/Nox2_TRN, 1,909 bp) and used as a specific probe in the following hybridization reaction (expected sizes: CgM2 1,763 bp, ΔCgnox2 3,572 bp, supplementary Figure S1). Re-integration of *Cgnox2* into the ΔCgnox2 deletion strain was tested using the primer pair nox2_P_comp_fw and nox2_eGFP_YR_rv (2,892 bp) in a PCR approach (supplementary Figure S2).

### 2.1.3 Chemotropic assay

Chemotropic growth towards different chemoattractants, maize root exudate (MRE), dihydrotanshinone I (DHT, Sigma Aldrich), and peroxidase from horseradish (HRP, Sigma Aldrich), was quantified after 6 h of incubation on agar in a 3D printed device as described previously (Schunke et al., 2020, Groth et al., 2021). The difference between attraction and non-stimulation was calculated shown with the calculated chemotropic index (Turra et al., 2015). Root exudate of maize plant was generated like described previously (Rudolph et al., 2024).

### 2.1.4 Analysis of leaf infection

The ability of *C. graminicola* asexual spores to penetrate maize plant material was analyzed on the second leaves of the *Z. mays* cultivar ‘Mikado’ (KWS SAAT SE, Einbeck, Germany). Unless otherwise stated, incubation of plants was performed in a PK 520 WLED plant chamber (Poly Klima Climatic Growth System, Freising, Germany) using a day/night cycle of 12 h 26°C/12 h 18°C. The strains CgM2, ΔCnox2, and ΔCgnox2::Cgnox2 were used for the infection experiments. Oval and falcate conidia were adjusted to 10^5^ spores per ml in 0.01% Tween. The second lowest maze leaves were fixated on wet blotting paper (BF2 580x 600mm, Sartorius, Göttingen, Germany) in square Petri dishes (82.9923.422, Sarstedt, Nümbrecht, Germany). Drops of 10 µl of the spore solution were added on top of the leaves. Analysis of symptom development was done after 5 days of incubation at 23°C with a rating of four different categories (no symptoms, minor symptoms, symptoms, severe symptoms) as described earlier (Nordzieke et al., 2019b). At least 40 individual spots were rated for fungal infection and mock infections (10 µl droplets of 0.01% Tween 20).

### 2.1.5 Analysis of root infection outgoing of spore-enriched vermiculite

The natural root infection outgoing from *C. graminicola* conidia present in growth substrated was simulated (Rudolph et al., 2024). Therefore, maize seeds were planted in 40 g of vermiculite (Vermiculite Palabora, grain size 2-3 mm, Isola Vermiculite GmbH, Sprockhövel, Germany) enriched with oval conidia of CgM2, ΔCgnox2 and ΔCgnox2::Cgnox2 in a concentration of 7.5 × 10^4^ x ml^-1^. As a mock control, seeds were sown in vermiculite mixed with water. Pots were sealed in disposal plastic bags (Sarstedt, Nümbrecht, Germany) to ensure high humidity. 21 dpi, length and biomass of the above-ground plant were determined.

### 2.1.6 Microscopy

Visualization of fungal structures on leaves was performed with light (differential interference contrast (DIC)) microscopy with the Axiolmager M1 microscope (Zeiss, Jena, Germany). The Photometrix coolSNAP HQ camera (Roper Scientific, Photometrics, Tucson, AZ, USA) was used to capture images. For image processing the ZEISS ZEN software (version 2.3, Zeiss) was used. For better visibility of penetration structures, leaf infection experiments were stopped after 3 d and the corresponding leaves stained with chlorazol Black E (Brachmann et al., 2001).

### 2.1.7 Statistics

For all experiments in this study, the T-test for unequal variances was used (Ruxton, 2006).

## 3 Results

### 3.1.1 Oval conidia of *Colletotrichum graminicola* are attracted by peroxidases

Tomato root exudated class III peroxidases (Prx) attracting *F. oxysporum* f. sp*. lycopersici* can be replaced by commercially available horseradish peroxidase (HRP) sharing a 37–39% amino acid similarity (Turra et al., 2015). In this study, the attraction of *C. graminicola* oval conidia to HRP was analyzed (Fig 1). Our data show that the amount of germlings able to re-direct growth to HRP is dose-dependent. The attraction of germlings peaked at an HRP gradient originating from 4 µM, whereas increasing concentrations did not provoke a redirection of germlings. However, comparable high doses of 128 µM HRP are again attracting germlings of the maize pathogen. Intriguingly, the resulting dose-response curve is highly similar to experimental data of *F. oxysporum* (Nordzieke et al., 2019a), raising the question whether molecular components required for Prx sensing might be conserved in *C. graminicola* as well.

**Figure 1.**
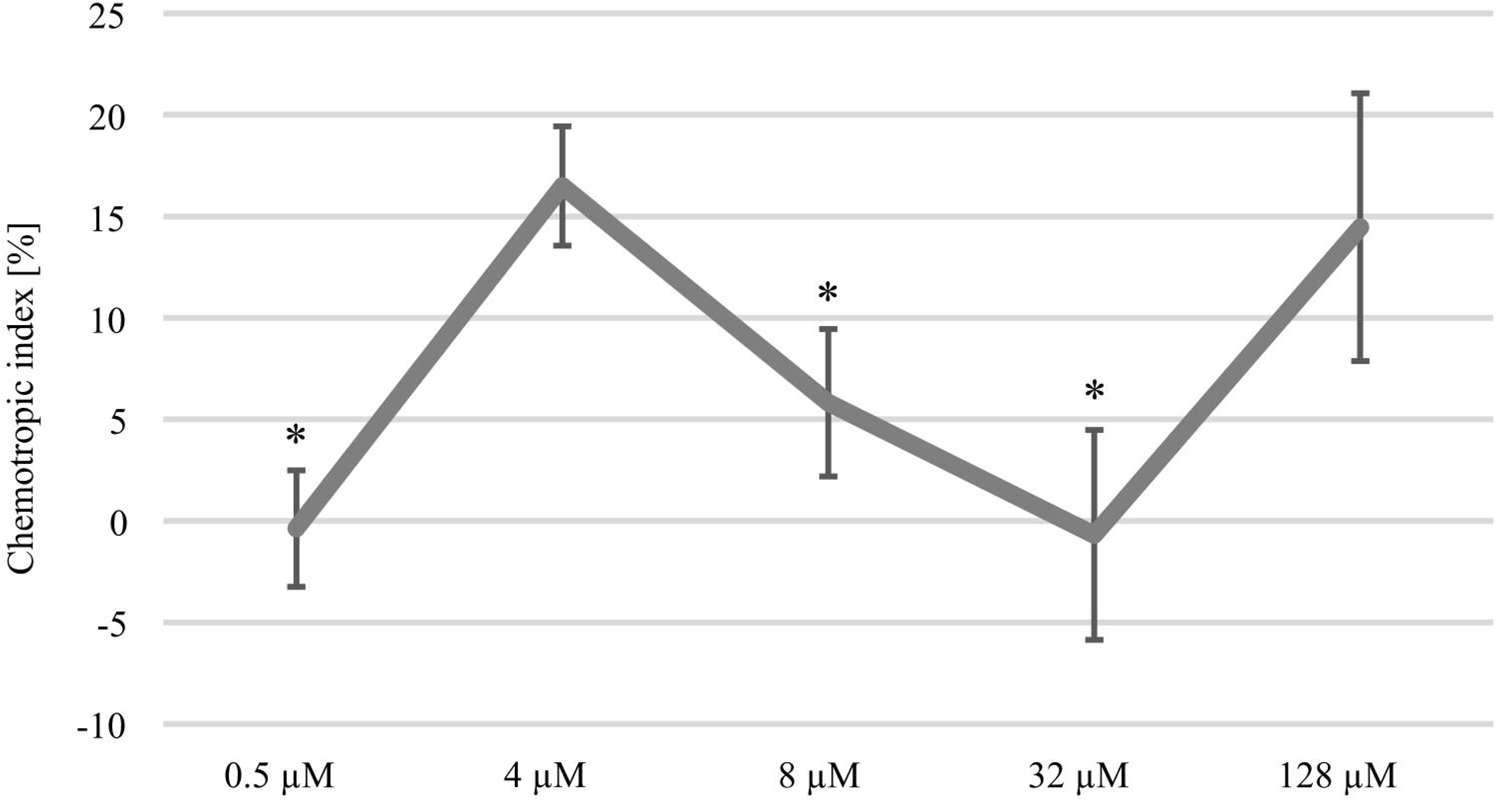
Dose-response curve to horse radish peroxidase (HRP). Chemotropic index displaying the attraction of CgM2 oval conidia by different concentrations of HRP. Redirection of growth was estimated after incubation for 6 h in a 3D printed device developed to analyze directed growth responses (Schunke et al., 2020). Error bars represent SD calculated from n ≥ 4 experiments, * p < 0.05, calculated with a two-tailed *t*-test.

### 3.1.2 Deletion of *Cgnox2* impairs the functionality of melanized penetration structures

Homologs of *Cgnox2*, the catalytic subunit of fungal Nox2 complexes, are responsible for plant-derived Prx sensing by *F. oxysporum* (Nordzieke et al., 2019a) and contribute to proper penetration structure function in several ascomycetes. To investigate the impact of *CgNox2* on the development of maize anthracnose, a deletion mutant of the corresponding gene was generated and analyzed regarding its impact on leaf and root infection processes.

Dependent on falcate or oval conidia inoculum, *C. graminicola* produces melanized penetration structures directly from spores (appressoria) or outgoing from fungal hyphae (hyphopodia), respectively (Nordzieke et al., 2019b). To assess the impact of *Cgnox2* deletion on the functionality of both infection structures, leaf infection experiments were conducted. Symptom rating after 5 days of inoculation revealed that independent of the spore type tested, ΔCgnox2 strains did not cause anthracnose lesions. This phenotype was fully converted in experiments using inocula of ΔCgnox2::Cgnox2 oval or falcate conidia (Fig 2 A and B). When we monitored fungal development in planta 3 dpi, melanized penetration structures were visible for all strains and spore types tested, indicating that the formation of appressoria and hyphopodia is not hampered by *Cgnox2* deletion. Detailed imaging of chlorazol-stained leaves revealed that primary hyphae are visible *in planta* in CgM2 and ΔCgnox2::Cgnox2 experiments. In experiments using ΔCgnox2 inocula, we were unable to monitor any leaf-colonizing hyphae, indicating that *Cgnox2* is required for proper penetration outgoing from appressoria and hyphopodia (Fig 2 C).

**Figure 2.**
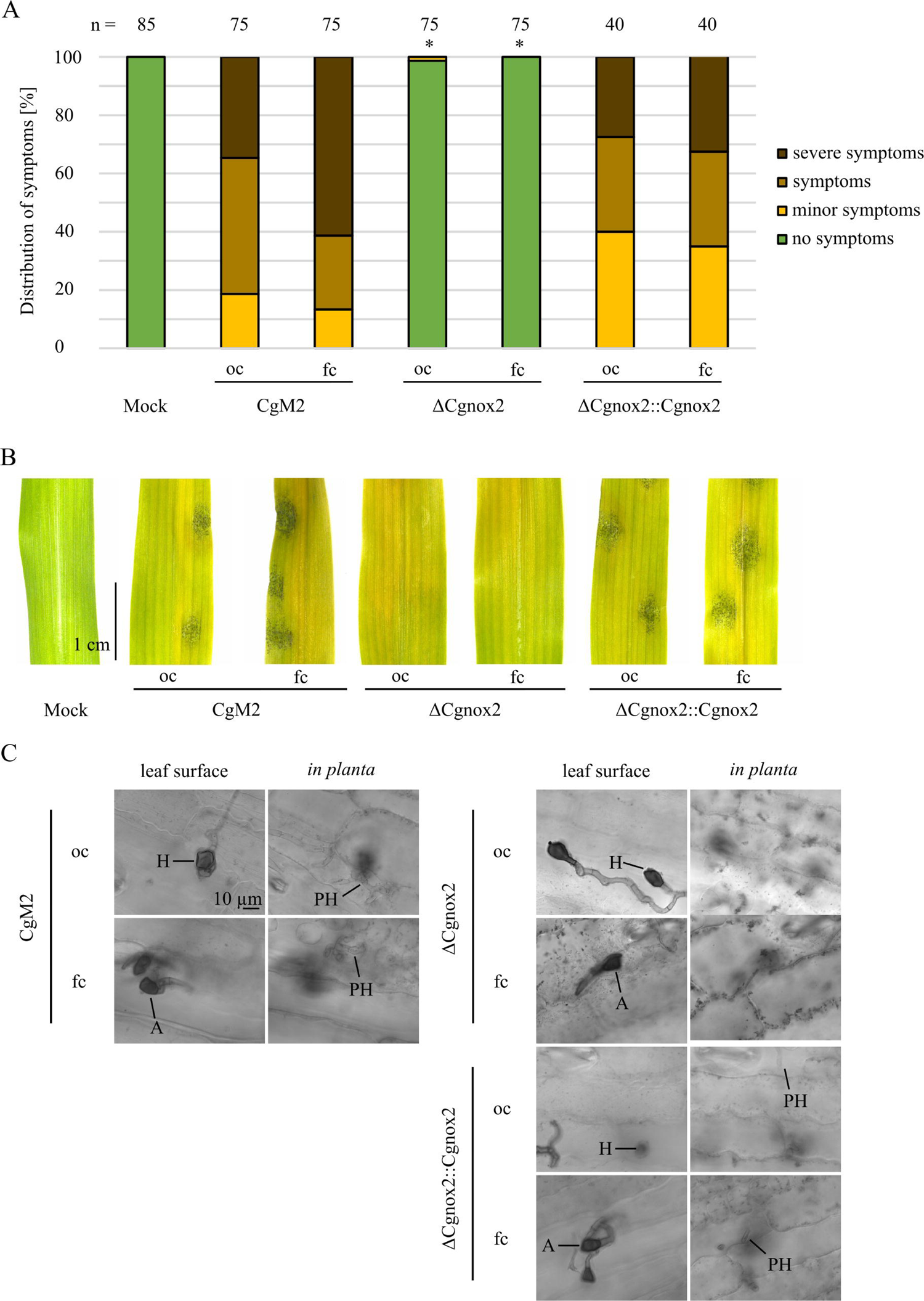
Symptom development on leaves provoked by different *C. graminicola* strains. **(A-C)** Infection of 16-day old maize leaves with oval and falcate conidia of the wild type CgM2, the mutant strain ΔCgnox2 and complemented strain ΔCgnox2::Cgnox2. 10^3^ spores were applied per infection site on the leaves. **(A-B)** Evaluation after 5 dpi. Symptoms are classified into four categories (no symptoms, minor symptoms, symptoms, severe symptoms) as described before (Nordzieke et al., 2019b). Infection spots evaluated n≥40. * p < 0.05, calculated with two-tailed *t*-test in comparison to CgM2 infected leaves. **(A)** Quantification of symptoms in percent. **(B)** Representative pictures of mock-treated and conidia-infected leaves. Size bar = 1 cm. **(C)** Monitoring of leaf infection sites 3 dpi. After staining with chlorazol Black E, cross-sections were taken, Size bar = 10 µm, H = hyphopodia, A = appressoria, PH = primary hyphae.

### 3.1.3 CgNox2 is dispensable for maize root infection and diterpenoid sensing

Next, we analyzed a probable involvement of CgNox2 on the infection of roots and sensing of maize root exudate and diterpenoids. For root infection experiments, we mixed oval conidia of different *C. graminicola* strains with vermiculite and inseminated the contaminated soil with maize seeds. This experimental setup mimics the natural infection situation in the field in which soil-borne fungal pathogens have to grow towards roots of their host prior to infection (Rudolph et al., 2024). As depicted in Fig 3 A and B, the *C. graminicola* wildtype CgM2 is able to stunt the above-ground tissue of growing maize plants. Likewise, the *Cgnox2* deletion strain showed a strong stunting phenotype, indicating that CgNox2 is dispensable for root sensing and its infection. These finding are supported by experiments in which the amount of germlings attracted by maize root exudate (MRE) and the diterpenoid dihydrotanshinone I (DHT) was analyzed (Fig 3 C). *C. graminicola* wildtype and ΔCgnox2 germlings show a strong chemotropic response to MRE as well as DHT, indicating that *Cgnox2* is not required for the sensing of the applied attractants.

**Figure 3.**
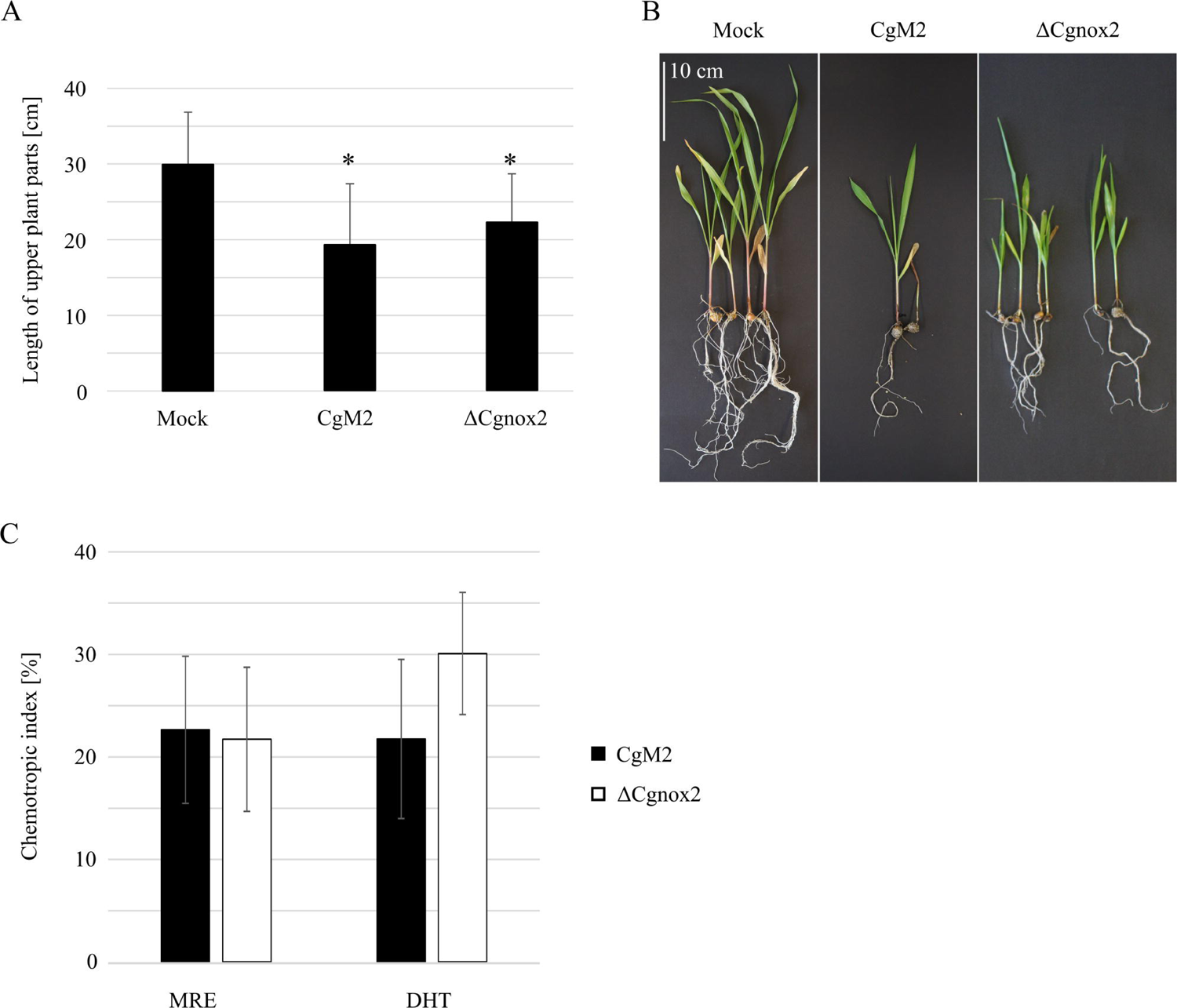
Relevance of CgNox2 for maize root infection. **(A-B)** Symptom development of plants co-incubated with oval conidia of CgM2 and ΔCgnox2. **(A)** Quantification of length of above-ground plant parts. The error bars represent the standard deviation from ≥ 6 plants, * p < 0.05, calculcated with two-tailed *t*-tests. **(B)** Representative depiction of the development of infected plants, scale bar = 10 cm. **(C)** Chemotropic index of CgM2 and ΔCgnox2 oval conidia facing gradients of maize root exudates (MRE) and the diterpenoid DHT, n ≥ 3.

### 3.1.4 Conserved pathway components mediate the perception of peroxidases and diterpenoids

The fungal pheromone receptors Ste2 and Ste3 of *Fusarium* species function in recognizing Nox2-activated peroxidase, inducing the phosphorylation cascade of the downstream CWI MAPK pathway (Turra et al., 2015, Sharma et al., 2022, Nordzieke et al., 2019a). To analyze whether a conserved Prx sensing machinery exists in *C. graminicola*, we confronted different deletion strains with the peaking HRP concentrations of 4 and 128 µM (Fig 1). Similar to findings in *F. oxysporum*, ΔCgnox2 germlings were unable to redirect growth to 4 µM HRP, but fully able to recognize 128 µM (Fig 4 A). A deletion mutant of the a-pheromone receptor gene *Cgste3* (Rudolph et al., 2024), however, was not attracted by both HRP concentrations applied. To investigate a probable requirement for CWI MAPK components, we further employed the role of *Cgso* (Nordzieke, 2022), encoding for a scaffolding protein of this pathway, for Prx sensing. Similarly, to ΔCgste3, the applied HRP gradients did not elicit a chemotropic growth response in ΔCgso. Attraction was fully restored in the complementation strains ΔCgnox2::Cgnox2, ΔCgste3::Cgste3, and ΔCgso::Cgso, indicating that central molecular components for Prx sensing are conserved among *F. oxysporum* and *C. graminicola*. For further characterization of diterpenoid sensing, we confronted *C. graminicola* wildtype, ΔCgste3, ΔCgste3::Cgste3, ΔCgso, and ΔCgso::Cgso germlings with MRE and DHT gradients (Fig 4 B). Similar to our observation for HRP sensing, the *Cgso* deletion mutant was unable to sense those molecules. Together, these results indicate that the sensing of chemically very different root-exudated molecules is routed via identical molecular pathways and is conserved among different fungal root pathogens (Fig 4 C).

**Figure 4.**
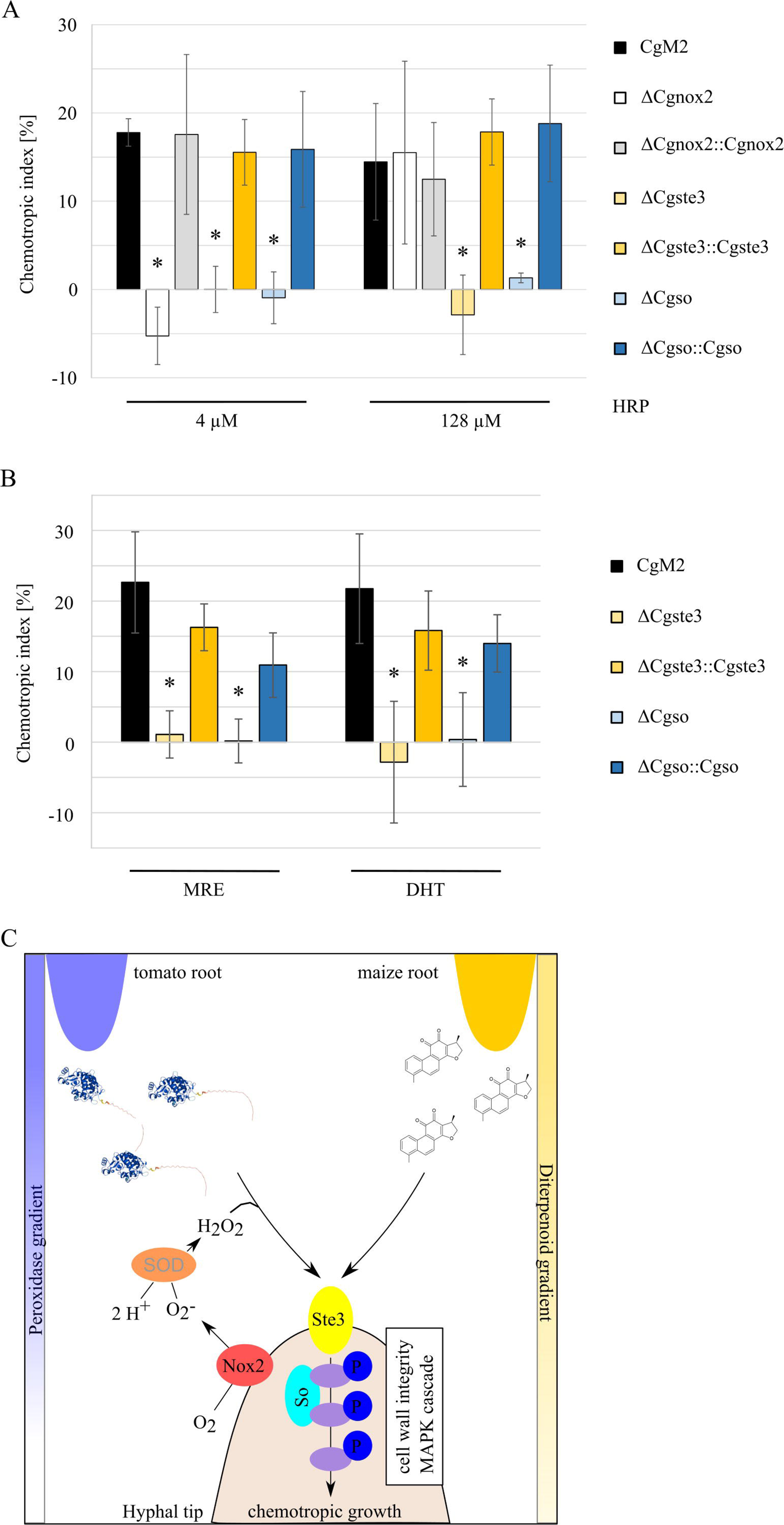
Signaling pathway components required for recognition of diterpenoids and peroxidases. **(A+B)** *C. graminicola* deletion mutants facing gradients of 4 and 128 µM (**A**) or maize root exudate (MRE) and the diterpenoid dihydrotanshione I (DHT, **B**) as indicated. (**C**) Model of conserved molecular pathways leading to growth re-direction due to peroxidase or diterpenoid gradient sensing, Nox2 = NADPH oxidase complex 2, Ste3 = a-pheromone receptor, So = scaffold protein of the cell wall integrity MAPK module, SOD = extracellular superoxide dismutase. Molecular factors investigated in this study or included from literature are written in black or grey letters, respectively. Molecule structure of peroxidase was generated by AlphaFold 3 (HRP_22489.1), chemical structure of DHT was drawn with ACD/ChemSketch.

## 4 Discussion

The rhizosphere is an environment full of different compounds such as nutrients and plant-based metabolites to attract or kill microbes (Vives-Peris et al., 2020). As numerous studies showed, plant exudates have a central role in shaping the root environment and are constantly adapted to fulfil the needs of the plant (Khorassani et al., 2011, Chaparro et al., 2013, Cheng and Cheng, 2015, Guerrieri et al., 2002). Knowing this changeable environment is crucial for plant-interacting fungi and a prerequisite to find their hosts and to avoid hazardous environments. Increasing evidence exist that fungal root pathogens have explored a possibility to hijack plant defense mechanisms to identify a close-by host (Turra et al., 2015, Vangalis et al., 2023, Sridhar et al., 2020, Ramaswe et al., 2024). In *Fusarium* and *Verticillum* species interacting with tomato plants, class III peroxidases (Prx) secreted from host roots induce a redirection of germling growth towards those plant defense molecules, instead of being repelled (Turra et al., 2015, Vangalis et al., 2023). Recently, we reported a similar growth response to be induced by plant-derived diterpenoids, a further known class of plant defense molecules (Rudolph et al., 2024, Mafu et al., 2018). In this study, we provide evidence that independent for their relevance for host recognition, *C. graminicola* is able to sense and react to Prx as well as ditpernoids. Several molecular factors like the a-pheromone receptor CgSte3 and the Cell Wall Integrity MAPK scaffold protein CgSo are required for the sensing of Prx and diterpenoids, despite the very different chemical properties of these molecules. In contrast, the activity of CgNox2 is specific for the recognition of Prx, but dispensable for diterpenoid sensing.

Prx take part in several plant defense processes, including cell wall enforcement, auxin metabolism, phytoalexin synthesis and the generation of reactive oxygen and nitrogen species (ROS, RNS) (Almagro et al., 2009). Despite of being defense enzymes, Prx are constantly expressed to a basal level, but their generation is accelerated in the presence of plant pathogenic fungi and bacteria (Sasaki et al., 2004, Young et al., 1995, Lavania et al., 2006, De-la-Pena et al., 2008). Several products of Prx activity like ROS, RNS and phytoalexins are well studied for their negative impact on membranes, resulting in lipid oxidations, alteration of the activity of membrane associated enzymes, impairment of membrane fluidity and increase of membrane permeability (Endale et al., 2023, Su et al., 2019, Di Meo et al., 2016, Jeandet et al., 2023). Diterpenoids are C20 compounds based on four isoprene (C5H8) units, which form a large and structurally diverse class of natural products found in plants, animals and fungi (Hanson, 2009, De Sousa et al., 2018). The antifungal activity against several maize pathogenic species including *B. cinerea* or *Rhizopus microsporus* was reported (Mendoza et al., 2009, Schmelz et al., 2011). As the products of Prx activity, diterpenoids can interact with membranes in various ways. For several diterpenoid molecules with antimicrobial functions, damaging interactions with membranes were reported (Saha et al., 2022, De Sousa et al., 2018) as well as their potential to induce ROS formation (Sun et al., 2021).

The a- and α-pheromone receptors Ste3 and Ste2 were first identified in yeast for their role in the recognition of the vice versa pheromones during mating (Nakayama et al., 1985, Jenness et al., 1986, Hagen and Sprague Jr, 1984). In filamentous fungi, the homologous G-protein coupled receptors (GPCRs) have acquired additional functions, including regulation of germination and the sensing of class III peroxidases and ditpernoids besides the sensing of pheromones (Vitale et al., 2019, Turra et al., 2015, Sharma et al., 2022, Rudolph et al., 2024). Together, these results raise the question, how a single GPCR respond to such a diverse set of molecules. GPCRs represent the largest class of signaling receptors known and are able to respond to various ligands, including protons, lipids, nutrients, and pheromones (Roth et al., 2017). After ligand recognition, GPCR activate downstream MAPK pathways, which in turn mediate the induction of various cellular responses (Xue et al., 2008, Braunsdorf et al., 2016). Besides the classical GPCR, there are several unconventional 7 transmembrane receptors. Adhesion GPCRs (aGPCRs) are characterized by an extended extracellular N-terminal region, which is able to bind several ligands each on a distinct protein fold (Araç et al., 2016). Other GPCRs can interact with other receptors types like protein tyrosine kinase receptors (PTKRs) and serine/threonine kinase receptors (S/TKRs) in a process termed transactivation (Schafer and Blaxall, 2017, Mohamed et al., 2019). Intriguingly, GPCRs are described to activate release of PTKR or S/TKR ligand upon activation, but also the activation of GPCRs by a primary activated PTKR or S/TKR is described (Kilpatrick and Hill, 2021). In this way, a GPCR can serve as a signaling hub for very different primary ligands. A central second messenger mediating transactivation are ROS, metabolites that are generated during peroxidase and diterpenoid plant defense responses.

Taken together, our results show that the 7 transmembrane G-protein coupled receptor (GPCR) CgSte3 is central for the sensing of chemically very different plant molecules as class III peroxidases and diterpenoids in the anthracnose fungus *Colletotrichum graminicola*. Upon activation, CgSte3 induces signaling via the downstream Cell Wall Integrity Mitogen Activated Protein Kinase pathway, resulting in a directed growth response of the plant pathogen towards a gradient of defense molecules, thus hitchhiking the original plant defense response. Upstream of CgSte3, we identified the NADPH oxidase CgNox2 as a specific factor for peroxidase sensing, which is dispensable for the perception of diterpenoids. The detailed molecular processes enabling a single GPCR to sense such chemically distinct molecules have to be revealed in future investigations.

## 5 Conclusions

This study explores the molecular mechanisms by which *Colletotrichum graminicola* interacts with plant root-secreted defense molecules. The 7-transmembrane G-protein coupled receptor (GPCR) CgSte3 is identified as crucial for sensing maize root exudates, including class III peroxidases and diterpenoids. Activation of CgSte3 triggers the Cell Wall Integrity Mitogen-Activated Protein Kinase (CWI MAPK) pathway, directing fungal growth towards these plant defense compounds. The NADPH oxidase CgNox2 is essential for peroxidase detection, highlighting a specific sensing mechanism. These findings reveal that CgSte3 and CWI MAPK pathways are central to *C. graminicola’s* ability to exploit maize defenses, offering promising targets for controlling maize anthracnose. Future research should focus on understanding how CgSte3 detects diverse signals, which could lead to innovative disease management strategies.

## 6 Data availability statement

Genetically engineered organisms and plasmids of this study are available from the corresponding author upon reasonable request.

## 7 Author contributions

AYR, CS and DEN conducted laboratory work, AYR and DEN wrote the manuscript, AYR, CS and DEN reviewed and edited the manuscript, DEN acquired funding and supervised the project.

## 9 Acknowledgements

We thank Gabriele Beyer and Gertrud Stahlhut for excellent technical assistance. The generative AI ChatGPT Version 3.5 was used to check this manuscript for grammar and typing errors.

## Supplementary Material

### 1 Supplementary Figures and Tables

#### 1.1 Supplementary Figure

**Figure S1.**
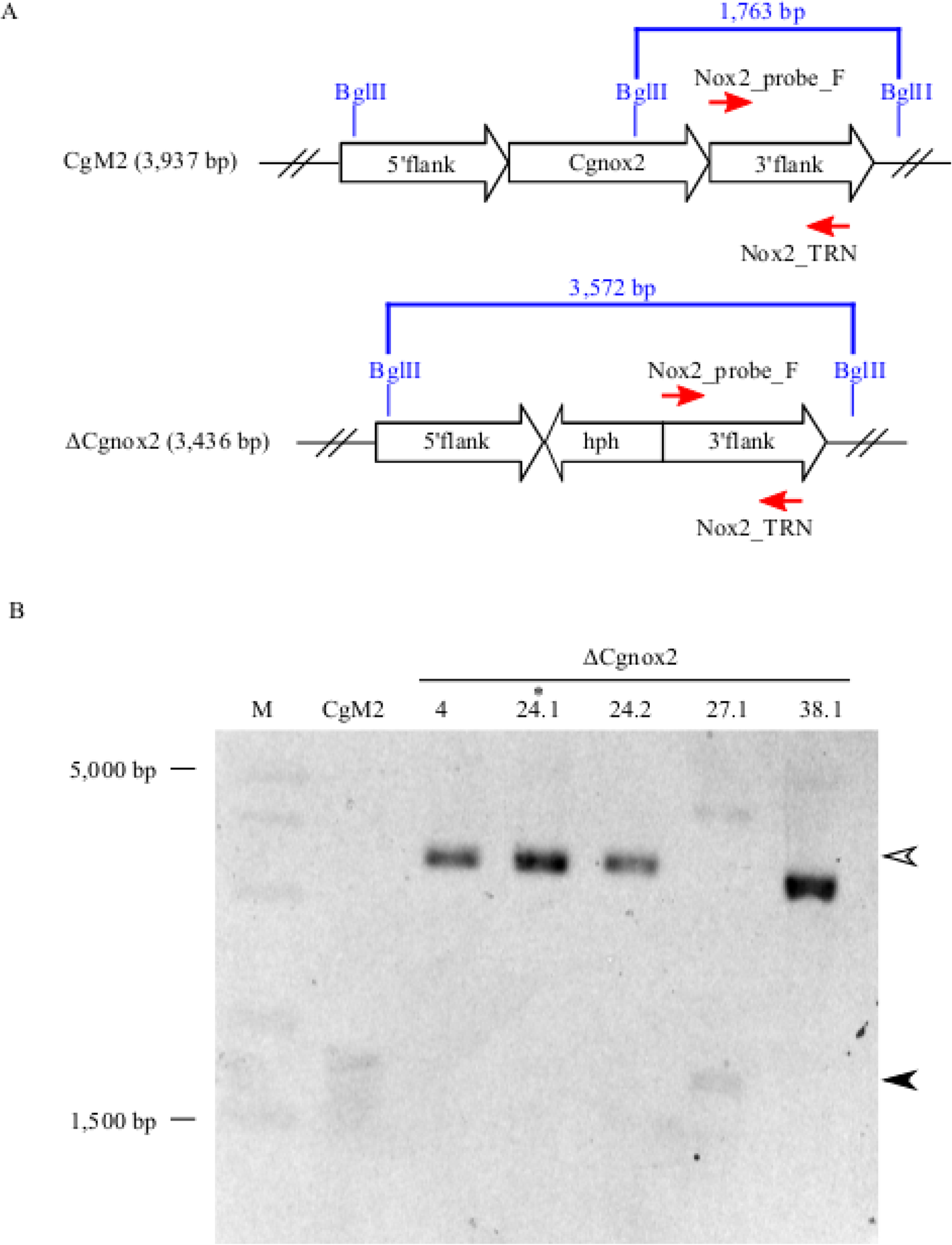
Generation of a *Cgnox2* deletion strain in *C. graminicola*. **(A)** Depiction of the Southern blot strategy. Genomic loci of *Cgnox2* in the wildtype strain CgM2 and the deletion strain. Primer binding sites for amplification of the 3′flank region for the probe. Red arrows indicate the amplification direction. Recognition sites of restriction enzyme *Bgl*II and the resulting band sizes are indicated in blue. **(B)** Southern blot hybridization for verification of homologous integration of the hph-resistance cassette into the *Cgnox2* locus. The strains used for phenotypic characterization are written in bold letters. The single spore isolate T24.1 (asterisk) was used for complementation.

**Figure S2.**
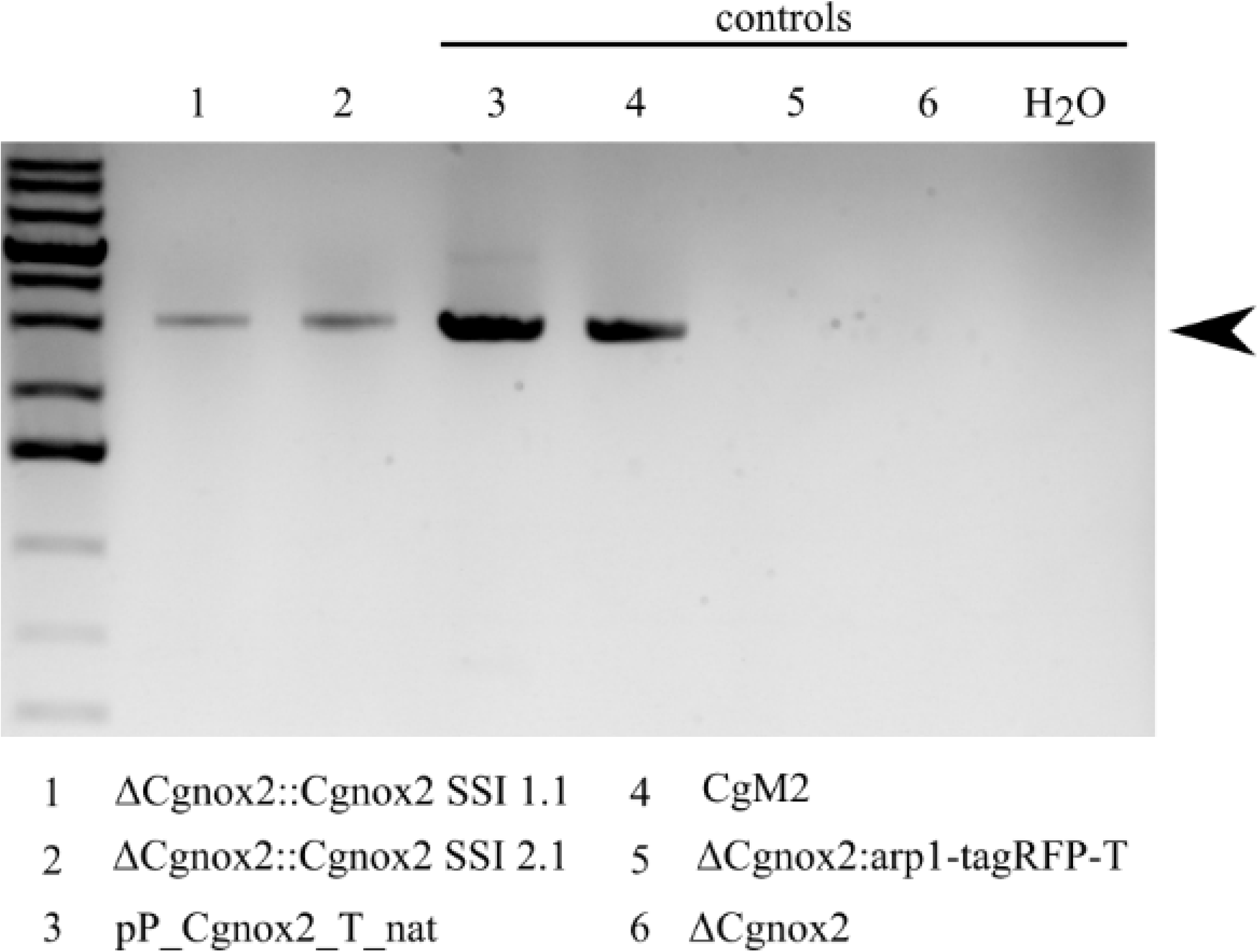
Verification of ΔCgnox2::Cgnox2. Amplification with the oligonucleotides nox2_P_comp_fw and noxB_eGFP_YR_rv for verification of ΔCgnox2::Cgnox2 single spore isolates T1.1 and T2.1. The strain used for phenotypic characterization is written in bold letters. The correct band size of 2,892 base pairs is indicated with a black arrow. Controls 1 and 2 have the expected bands, while negative controls 3 and 4 show no amplification.

#### 1.2 Supplementary Tables

**Table S1.**
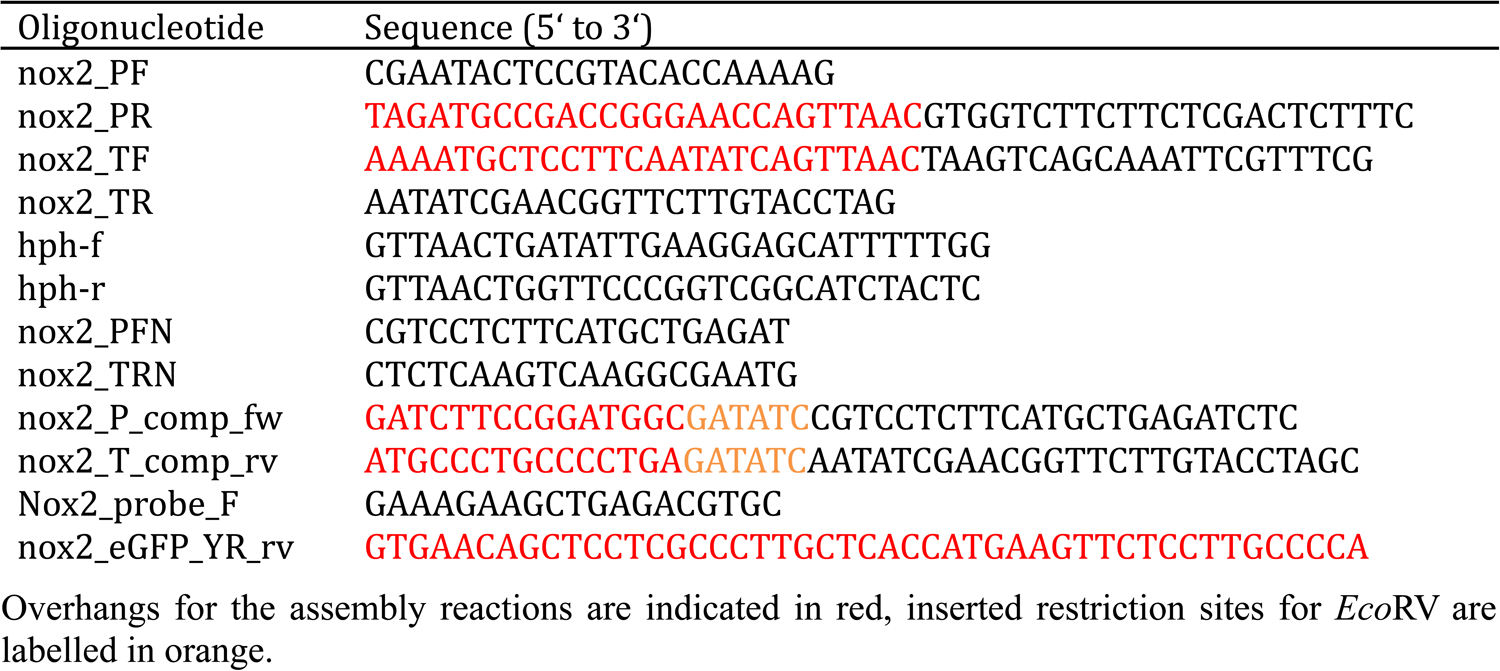
Oligonucleotides used in this study.

**Table S2.**
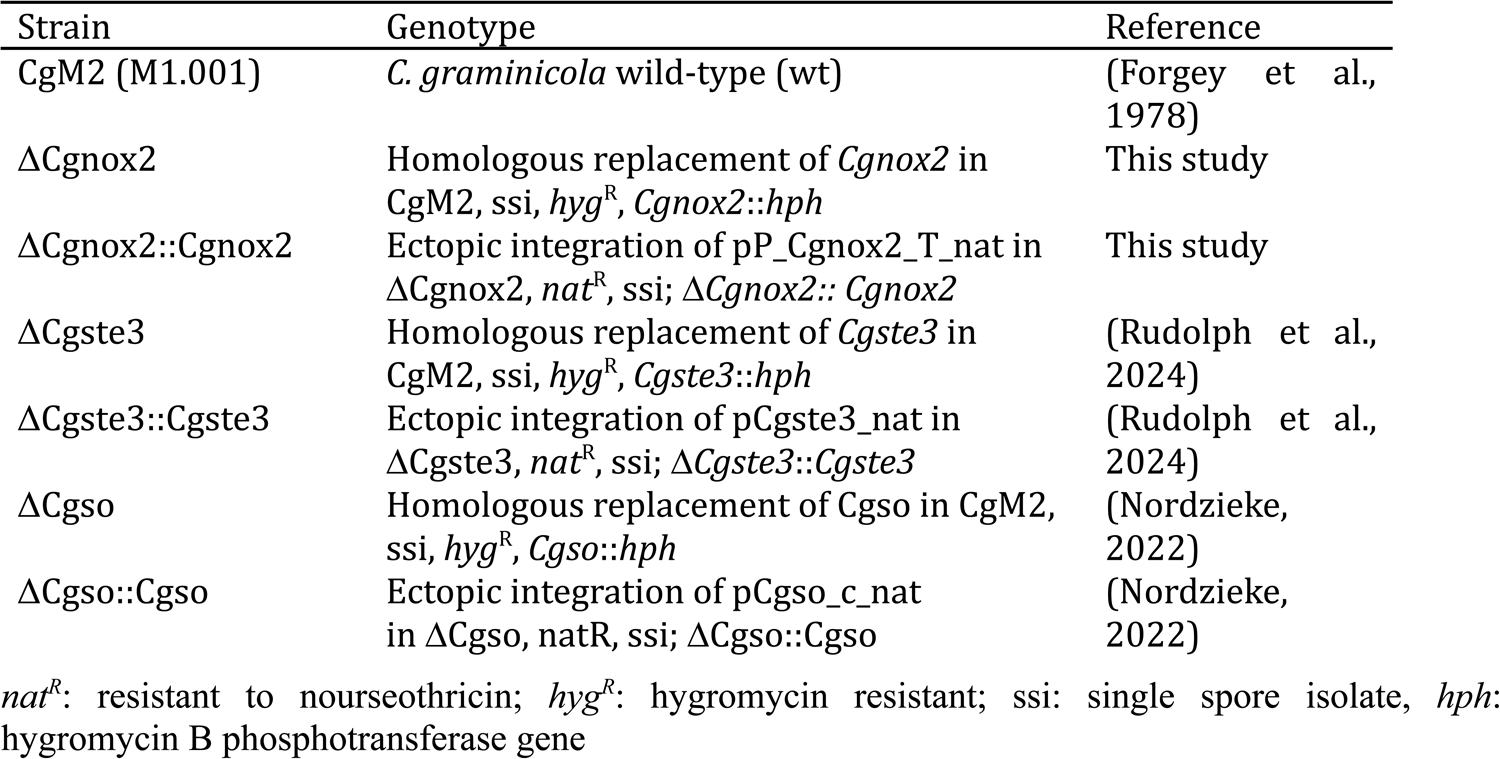
*Colletotrichum graminicola* strains used in this study.

**Table S3.** Plasmids used in this study.

| Plasmid | Features | Reference |
| --- | --- | --- |
| pCgnox2_KO | 5' *Cgnox2*::*hph*::3' *Cgnox2*, *hyg*<sup>R</sup>, *amp*<sup>R</sup> | This study |
| pCgnox2_nat | 5' *Cgnox2*::*Cgnox2*::3' *Cgnox2*, *nat*<sup>R</sup>, *amp*<sup>R</sup> | This study |
| pJet1.2 | *amp*<sup>R</sup> | ThermoFisher Scientific |
| pJet_nat | *nat*<sup>R</sup>, *amp*<sup>R</sup> | (Nordzieke, 2022) |

